# Microtubule Defect Involved in ‘Mitophagy Resistance’ Under Subacute Oxidative Stress - Potential Mechanism for Cellular Inflammation

**DOI:** 10.1101/2020.02.26.966234

**Authors:** Takahiko Tamura, Nobuo Yasuda, Tomoharu Shakuo, Aki Kashiwagi, Jeevendra A. J. Martyn, Masataka Yokoyama, Shingo Yasuhara

## Abstract

**Introduction:** Oxidative stress is considered an essential mechanism in ICU-acquired weakness. The roles of oxidative stress in autophagy/mitophagy dysfunction remains elusive. Microtubule serves as an essential guide rail for auto/mitophagosome trafficking required for proper maturation of auto/mitophagosomes in normal circumstances, and microtubules network formation is regulated by signal transduction mechanisms involving Akt, GSK3β, and the microtubule plus-end tracking molecule, EB1. We have investigated (1) whether oxidative stress affects this pathway, leading to the defective mitophagy response, and (2) whether trehalose, an auto/mitophagy modulator, can ameliorate these pathological conditions.

**Methods:** By stably transfecting markers for auto/mitophagy or MT synthesis, we have established a few new C2C12 myocyte cell lines, expressing, GFP-LC3, EB1-GFP, and/or tandem-fluorescence LC3 (tfLC3). To monitor microtubule network, the cells were stained by SiR-tubulin. The cells were cultured in the presence or absence of oxidative stress by hydrogen peroxide (H2O2) and treated with or without trehalose. The response of mitophagy parameters including vesicle motion and the maturation status was monitored by stimulating the cells with carbonyl cyanide m-chlorophenyl hydrazone (CCCP), an established mitophagy inducer, under a time-lapse confocal microscopy. Signal transduction mechanisms linking mitophagy to microtubule formation was analyzed by Western Blotting against Akt and GSK3β.

**Results:** Cells under the oxidative stress, showed abolished MT network formation, decreased microtubule synthesis by EB1, and a decrease in CCCP-invoked response of mitophagosome motion, perturbed mitophagosome maturation, and increased superoxide production. Signal resistance of Akt/GSK3β pathway to mitophagic stimulation, was documented. Trehalose treatment reversed signal resistance, diminished MT synthesis, ameliorated the disturbed MT network, and improved maturation defects, suppressing the production of superoxide.

**Conclusions:** Oxidative stress decreases the response of mitophagy and abolishes microtubule network. Trehalose improves the synthetic ability of microtubule and normalized the disturbed microtubule network, resulting in the improvement of the perturbed mitophagosomes maturation under the oxidative stress.

## Introduction

Muscle wasting and muscle weakness is one of the major complications among many types of critical illnesses including sepsis, burn, and major trauma [1]. In critical illnesses, muscle wasting occurs mostly in chronic phase. More recently, however, ICU-acquired weakness (ICU-AW), which occurs rapidly in the critically ill patients, has acquired both scientific and clinical attention in the field of critical illness studies [2] and the onset of muscle wasting/dysfunction is regarded to develop at relatively earlier stage of the disease than previously considered. When the disease persists, the muscle wasting and weakness arising from these illnesses lead to prolonged mechanical ventilation [3], with increased morbidity and mortality [4]. Oxidative stress and mitochondria dysfunctions are considered one of the key mechanisms in sepsis or burn injury-induced organ dysfunctions, and in ICU-AW. [5, 6]

Mitochondrial dysfunction has been reported to occur concomitantly in many forms of muscle dysfunctions in critical illnesses. Perturbation of mitochondrial network integrity and their functions leads to systemic catabolism, affecting adenosine triphosphate (ATP) production, decreased organelle biogenesis, elevated proteolysis and often resulting in the loss of muscle size [2]. In addition to their roles in bioenergetic homeostasis, mitochondria are involved in various signal transduction. Dysregulation of these signals lead to cellular malfunction directly or indirectly. Damaged mitochondria release catastrophic mediators including reactive oxygen species (ROS), cytochrome-c, which can initiate the process of cell death and induce proteolysis [7]. Mitochondrial quality control is therefore essential not only for energy homeostasis but also in the determination of cell fate.

Autophagy is a cellular housekeeping system that mediates either bulk removal of cellular components or selective degradation of damaged organelles and protein aggregates [8]. Its defect or derangement in signals is associated with pathogenesis of various diseases such as cancer, neurodegeneration, metabolic diseases and multiple organ failure of critically illness [9]. Autophagy plays an essential role in the skeletal muscle system, maintaining muscle fiber integrity [10]. Dysregulation of autophagy [5, 11] has been associated with the development of ICU-AW, or muscle wasting syndrome in critical illnesses, but the precise pathophysiological mechanism has remained elusive. Particularly, damaged mitochondria are removed by a form of selective autophagic degradation, or mitophagy. Mitophagy thus plays essential roles of cellular and mitochondrial homeostasis, but the causative relationship with muscle wasting and mitophagy has not been fully clarified in critical illness-induced muscle dysfunctions or ICU-AW despite the reports of mitochondrial dysfunctions. [12]

The relationship between oxidative stress and auto/mitophagy has been studied intensively mostly with the conclusion that oxidative stress, mediated by ROS, activates auto/mitophagy [13, 14]. Given that auto/mitophagy are cell protective systems acquired by eukaryotes during evolution [8, 15], findings about ROS activating auto/mitophagy is rational. The cells need to adapt to acute stress by reinforcing auto/mitophagy. In this context, however, the effect of oxidative stress on the basal level of, (but not the responsiveness or competency of), auto/mitophagy, has been the focus of study [16]. The effect of prolonged oxidative stress on the ‘response’, or competency, of auto/mitophagy has not been investigated in detail. Clinically, it has been recognized that competency of various organs for stress adaptation is diminished in many chronic inflammatory diseases or in critical illnesses. [17, 18] In this setting, it can be advocated that examining the competency of, but not just the basal level of, auto/mitophagy, is critical to fully understand the pathological symptoms of many diseases. This study, therefore, focused on the analyses of the capability of cells for turning mitophagy flux, in response to mitophagy-inducing stimulation and investigated how oxidative stress affects this response. Herein, by focusing on the response of mitophagy, a new disease entity of compromised response to mitophagy-inducing stimuli (’resistance to mitophagic stimuli’, or operationally simplified as ‘mitophagy resistance’) will be discussed. Similar to insulin resistance in type 2 diabetes mellitus (T2DM) or in chronic illnesses, we have noted there is a pathological condition where signal response for mitophagy flux turnover is defective, thus yielding poor response to mitophagy-invoking stimuli and thus resulting in decompensated quality control of mitochondria and in diminished safety margin in the homeostasis of mitochondria. [19, 20]

Furthermore, despite the intensive studies about the relationship between ROS and auto/mitophagy, the molecular target of ROS in auto/mitophagy pathway remained somewhat elusive [16]. Previous studies tried to link ROS signal to pathways for activating the basal level of auto/mitophagy signals [13, 21], but studies for explaining the link between the ROS and the responsiveness of auto/mitophagy have been limited. Microtubule (MT)s are hollow cytoskeletal rods with a diameter of approximately 25nm, and their network is maintained by a dynamic equilibrium, via constantly undergoing assembly and disassembly to fulfill cellular needs. They function both to determine cell morphology and participate in a variety of cell locomotion, the intracellular transport of organelles, and the separation of chromosomes during mitosis. Accumulating numbers of recent studies demonstrated that MT serves as an essential guide rail for auto/mitophagosome vesicle trafficking and thus are the key component of auto/mitophagy pathway [22]. The roles of MTs in auto/mitophagy dysregulation, however, have less been investigated especially in the context of diseases. In this study, we analyzed whether MTs and its regulation are affected by oxidative stress and how it leads to the defective mitophagy response.

There have been attempts of therapeutic approaches by stimulating the autophagy in disease models where defects of autophagy and/or mitophagy is reported. Many of these approaches are aimed at activating the upstream autophagy initiating mechanisms. However, there has been no discussion regarding the validity of stimulating the upstream event where the downstream can be defective and thus the flux potentially be blocked due to the disease. The difficulty in such discussion derives partly due to the elusiveness of pathophysiology of auto/mitophagy defect in many diseases. In this study, after evaluating the flux blockade of mitophagy response, the efficacy of a non-conventional auto/mitophagy modulator is examined. Trehalose, a non-reducing disaccharide composed of two D-glucose units linked α-1,1, is present in many organisms, including bacteria, fungi, plants, yeast, and invertebrates including tardigrades and brine shrimps. These organisms use trehalose to augment their adaptation competency against extremely severe environmental conditions including frigidity, dehydration, starvation, and UV-radiation. [23, 24] Supplementation with trehalose also improves the survival of mammalian cells [25, 26]. Recent studies demonstrated that trehalose induces autophagy via mammalian (mechanistic) target of rapamycin (mTOR)-independent pathway [27, 28]. Some studies have suggested that trehalose helps maintain MT-forming function of tubulins [29]. By employing the cell-protective and autophagy-augmenting proficiency of trehalose, therapeutic studies have been proposed to treat several diseases in which autophagy plays an important role [9]. This study therefore examined whether normalizing the disturbed MTs with trehalose can prevent the mitophagy dysfunction under the subacute oxidative stress.

## Materials and Methods

### Ethics Statement and Animal Research

There is no animal research involved in the current study.

### Reagents, Transgene Plasmid Constructs, Antibodies

The reagents used in this study are; MitoSOX (Thermo Fisher Scientific), LysoTracker (Thermo Fisher Scientific), SiR-tubulin (Cytoskeleton), CCCP (Millipore Sigma), E64d (Millipore Sigma), pepstatin A (Millipore Sigma), Lipofectamine 3000 (Thermo Fisher Scientific), G418 (Millipore Sigma), Mammalian expression plasmids for GFP-LC3 (kindly provided by Dr. N. Mizushima), tandem-fluorescent LC3 (Addgene), GFP-EB1 (Addgene), antibodies against LC3 (Sigma), phospho-Akt (Ser473, Cell Signaling Technology, #9271), phospho-GSK3β (Ser9, Cell Signaling Technology, #9322), Anti-GAPDH (#9484 abcam, #2275-PC-100 Trevigen)

### Cell Culture and Staining

C2C12 murine myocytes were obtained from ATCC and maintained in DMEM (Gibco/Thermo Fisher Scientific) with 10% fetal bovine serum (ATCC) and 50U/mL penicillin-streptomycin (Gibco/Thermo Fisher Scientific). By stably transfecting auto/mitophagy markers in C2C12 myocyte, new muscle cell lines were established, expressing GFP-LC3 or EB1-GFP or tfLC3.

### Stable Transfection

Mammalian expression transgene constructs were transfected into C2C12 cells by lipofection using Lipofectamine 3000 according to the manufacturer’s instructions. After 24 hours of transfection, stable transfectants were selected using G418 selection for one week and fluorescently labeled cells were cloned according to the method described previously [30].

### Drug Treatment, Mitophagy Induction, and Oxidative stress

To impose subacute oxidative stress, myocytes were exposed to 500μM of hydrogen peroxide (H_2_O_2_) for 12-18 hours. To test the therapeutic efficacy of trehalose, the cells were treated with or without 100μM trehalose for 2 hours before the observation. To monitor the response of mitophagy, the cells were stimulated with 5μM carbonyl cyanide m-chlorophenyl hydrazone (CCCP), an established mitophagy inducer. To disrupt microtubule network, the cells were treated with 5μM colchicine for 1 hour.

### Cell Staining

To monitor lysosomes, the cells were stained by LysoTracker Blue. To observe microtubule, the C2C12 myocyte were stained by SiR-tubulin. To investigate the superoxide production from mitochondria, the C2C12 cells were stained by MitoSOX at 5μM.

### Image Data Analysis

For detailed time-lapse image analyses, C2C12 cells were observed using a confocal microscopy (Nikon Eclipse, A1R HD) with a 2 second interval, and the captured images were analyzed by Image J. To investigate the mechanism of the beneficial function of trehalose on MT, the synthesis speed of MT was monitored using EB1-GFP, the plus-end tracking reporter. To investigate the mitophagy response induced by CCCP, the motion and maturation of mitophagosome were monitored. The vesicle motion was measured by using the tracking plugins of ImageJ. The maturation of mitophagosome was defied by the ratio of RFP/GFP. To measure fluorescent light intensity, a wide-field fluorescent microscope (Nikon Eclipse 800) calibrated with a standard fluorescent beads (LinearFlow, Thermo Fisher)

### Cell Experiment Procedure for Western Blot

The C2C12 cells were cultured on 6cm dishes to 80-90% confluence. Cells were serum starved, treated with or without oxidative stress by using 500μM H_2_O_2_ for 12-18 hours. After that, the cells were treated with or without 100μM trehalose for 2 hours. The response of mitophagy was monitored by stimulating the cells with 5μM CCCP.

### Cell Homogenization

Cell samples were frozen under liquid nitrogen and homogenized in ice-cold homogenization buffer (50 mM HEPES-NaOH, pH 7.5, 150 mM NaCl, 2 mM EDTA, 1% Nonidet P-40, 10% glycerol, 10 mM sodium fluoride, 2 mM sodium vanadate, 1 mM phenylmethyl-sulfonyl fluoride, 10 mM sodium pyrophosphate, 10 μg/ml aprotinin, and 10 μg/ml leupeptin). The homogenized samples were centrifuged at 13000 rcf for 30 min. Aliquots of supernatant containing same amounts of protein, determined by BCA protein assay method.

### Western Blot

Cell homogenates containing equal amounts of protein were subjected to 10% SDS-PAGE after the addition of sample buffer and boiling for 3 min. After being transferred electrophoretically to nitrocellulose membrane (Li-COR, Lincoln, NE), the membranes were blocked in 5% skim milk for 1 h at room temperature, followed by incubation with primary antibodies for overnight at 4°C.

Anti-phospho-Akt (p-Akt), anti-phospho-GSK-3β (p-GSK-3β), anti-GAPDH (Cell Signaling, Beverly, MA), anti-LC3 (Sigma Aldrich, St Loius, MO) antibodies were used as primary antibodies. The anti-rabbit or -mouse IgG antibody was used as a secondary antibody at a dilution of 1:10,000. Bands of interest were scanned and quantified with the use of Odyssey CLx (Li-COR, Lincoln, NE).

### Statistical Analysis

Data are expressed as mean±standard deviation (SD) and analyzed with Student’s t-test for two-group comparison, one-way ANOVA and Tukey’s method for more than three-group comparison. A value of p<0.05 was considered significant.

## Results

### Subacute Oxidative Stress-Induced Perturbation in Mitophagy Response

To investigate the non-acute effect of oxidative stress on the responsiveness of mitophagy, C2C12 cells were treated with H_2_O_2_ for 16 hours. Cells were then stimulated by CCCP to induce mitophagy in the presence or absence of auto/mitophagy flux blockade with E64d and pepstatin A. Purified mitochondria fraction was analyzed for mitophagy induction by Western Blotting against LC3. As shown in Fig.1, H_2_O_2_ treatment elevated the basal level of mitophagy, but when CCCP-stimulated response of mitophagy was compared between controls (H_2_O_2_ -) and the oxidative stress group (H_2_O_2_ +), mitophagy flux was significantly blocked, indicating that prolonged exposure to oxidative stress confers signal resistance to mitophagic stimulation in C2C12 cells.

**Figure 1:**
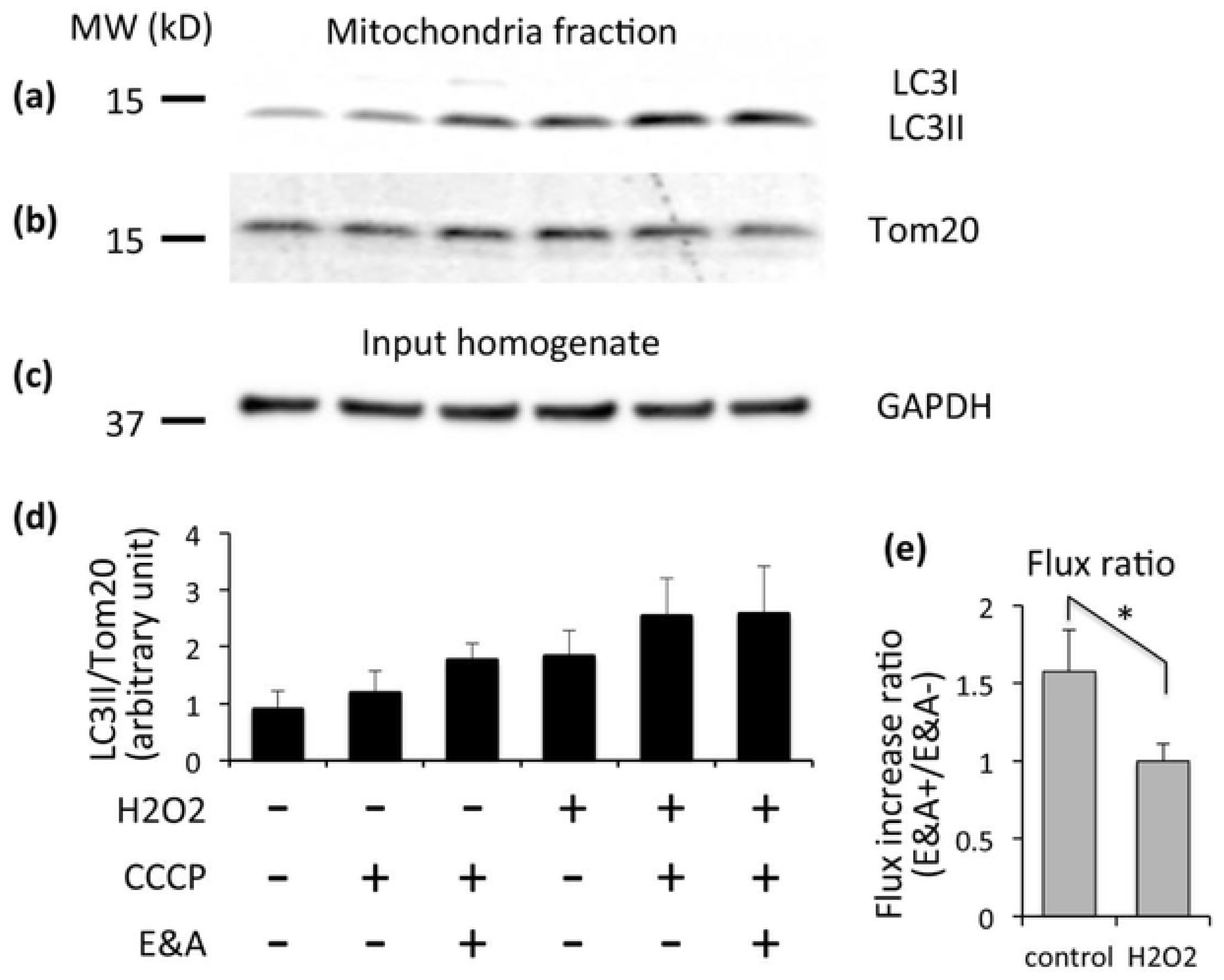
Mitophagy flux response to CCCP stimulation is diminished under oxidative stress. (a-d) Western blotting pictures either of the mitochondrial fraction (a, b) or the input of total homogenate (c) are shown. C2C12 myocytes were cultured in the absence (“H_2_O_2_-”) or presence (“H_2_O_2_+”) of hydrogen peroxide for 18 hours. Cells were stimulated by CCCP and the response of mitophagy was monitored with or without mitophagy flux blockade by E64d plus pepstatin A (“E&A-” vs. “E&A+”). Shown in the WB pictures are; (a) blotting against LC3, the auto/mitophagy marker in the mitochondrial fraction, (b) mitochondrial residual molecule, Tom20, as an internal control for equal mitochondrial protein load, and (c) GAPDH in the total homogenate as the input loading control for fractionation. (d) Quantification of the band intensity is shown as the ratio of LC3II/Tom20 in the mitochondrial fraction. Despite the tendency of increased basal level of mitophagy under oxidative stress without mitophagy stimulation, (H_2_O_2_+, CCCP-, E&A-), mitophagy flux is diminished (increment from E&A- to E&A+) under the oxidative stress (H_2_O_2_+), as compared to that in the control group (H_2_O_2_-). (e) The ratio of increase in the LC3II/Tom20 before and after the flux blockade is shown. *: p<0.05, N=4. Data are shown as average +/- standard deviation. Note that in the mitochondrial fraction for LC3 (a), LC3II form is predominantly detected, because this form mainly consists of the vesicle attached form.

### The Effect of Oxidative Stress on Mitophagy Motion and Its Maturation

To further investigate the mechanisms involved in the disturbed mitophagy response by CCCP under oxidative stress conditions, the movement of mitophagosome was monitored by time-lapse confocal microscopy, using C2C12 cells expressing GFP-LC3. In cells under the subacute oxidative stress, the movement of mitophagosomes invoked by CCCP decreased significantly than that in the control cells (Fig.2, Supplementary Video 1), consistent with the data for the decreased mitophagy flux under CCCP stimulation in Fig.1. Trehalose, however, ameliorated the once diminished movement of mitophagosomes under the subacute oxidative stress (Fig.2, Supplementary Video 1). The increment of mitophagosome motion in response to CCCP was significantly improved by trehalose treatment under the subacute oxidative stress (132 vs. 100%: with vs. without trehalose; p<0.05).

**Figure 2.**
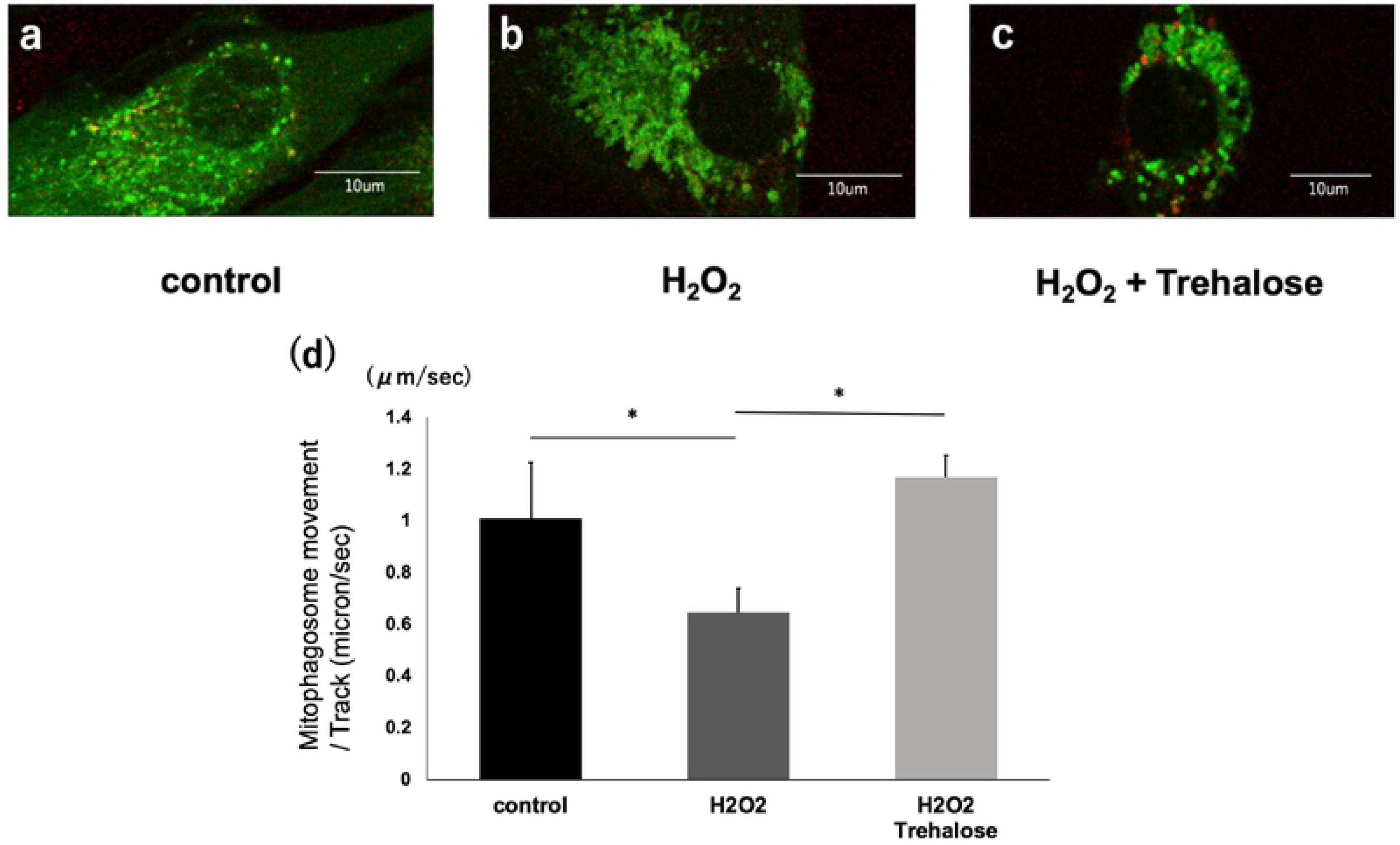
The mitophagy movement is diminished under oxidative stress. (Top) (a-c) Confocal microscopic images of C2C12 cells expressing GFP-LC3, co-stained with LysoTracker Blue. Pseudo colors are added with green for GFP-LC3, and red for LysoTracker Blue. (a) In controls, GFP-LC3 dots are fine granular and show vigorous movement. (b) In H_2_O_2_ treated cells, GFP-LC3 vesicles are swollen and the motions diminished. (c) Trehalose treatment partially reverts the size of GFP-LC3 dots and reactivates the vesicle motion. White scale bar at the right bottom corner of each image represents 10μm. (d) The movement of mitophagosomes was analyzed by ImageJ and shown as the average value of area with standard deviation (N=3-8). *:p<0.05 by ANOVA with Tukey’s comparison. For video information, refer to Supplementary Video 1. Supplementary Video 1: Time-lapse confocal video image of C2C12 myocytes expressing GFP-LC3, stained by LysoTracker Blue. Live video information for Fig.2 is shown here. Cells are incubated in the presence or absence of H_2_O_2_ with or without trehalose treatment. In controls (H_2_O_2_ -, trehalose -, far left), GFP-LC3 dots are fine granular and show vigorous movement. In H_2_O_2_ treated cells without treatment (H_2_O_2_ +, trehalose -, middle), GFP-LC3 vesicles are swollen and the motions markedly diminished. Trehalose treatment (H_2_O_2_ +, trehalose +, far right) partially reverts the size of GFP-LC3 dots and reactivates the vesicle motion. Yellowish dots represent the mature vesicles after LC3-GFP dot (green) fuses with lysosomes (red), and also shows similar pattern of motion changes. In this video, pseudo colors are added with green for GFP-LC3, and red for LysoTracker Blue.

To investigate the impact of decreased motion of mitophagosomes due to oxidative stress on the mitophagy flux, C2C12 cells expressing tandem-fluorescent LC3 (tfLC3) were monitored for analyzing the maturation status and thus flux of mitophagy after CCCP stimulation. As is shown in Fig.3, tfLC3-expressing cells under the oxidative stress harbored more yellow dots (GFP and RFP both positive) as compared to red-only predominant pattern in controls, demonstrating the inhibition of mitophagosomes maturation under oxidative stress conditions. Note that tfLC3 reporter provides fluorescent signal of both GFP and RFP and thus yield yellow signal when autophagosomes and mitophagosomes are premature. During the process of maturation, autophagosomes and mitophagosomes fuse with lysosomes, and due to the drop of intravesicular pH, GFP signal fades, resulting in red-dominant signal. This finding of defective maturation supports the decreased mitophagy flux under the oxidative stress (Fig.1). Trehalose treatment, however, improved ratio of yellow to red mitophagosomes, and ameliorated the defective maturation under the subacute oxidative stress. Next, to further decipher the mechanism of the beneficial effect of trehalose, colchicine, a drug that destabilizes microtubule, was added to the treatment. Colchicine abolished the effect of trehalose on mitophagy maturation (Fig.3), suggesting that beneficial effect of trehalose is likely mediated by its effect on normalization of microtubules.

**Figure 3.**
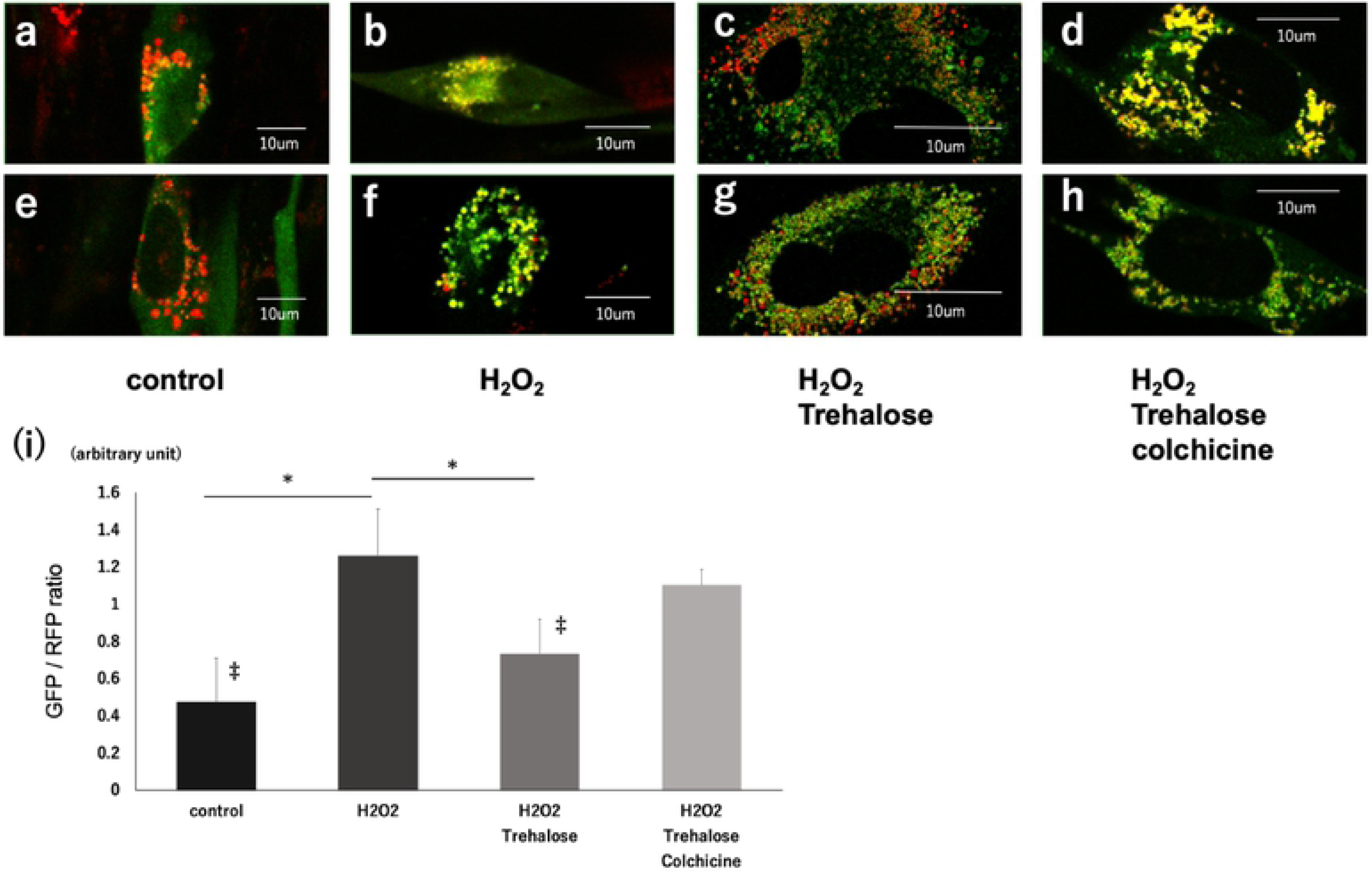
The mitophagosome maturation is disturbed under oxidative stress. (Top) (a-h) Confocal microscopic images of C2C12 cells expressing tf-LC3 are shown to document the effect of H_2_O_2_ and Trehalose on the maturation of mitophagosomes after CCCP stimulation (GFP Green, RFP; Red). (a,e) Control group, (b,f) H_2_O_2_ group, (c,g) H_2_O_2_ +trehalose group, (d,h) H_2_O_2_ +trehalose +colchicine group. Scale bar represents 10μm. (i) The maturation status of mitophagosomes was analyzed by ImageJ and shown as the average value of the ratio of areas of GFP/RFP with the standard deviation (N=12). *:p<0.05, ‡:p<0.05 vs colchicine group by ANOVA with Tukey’s comparison.

### The Effect of Oxidative Stress on Microtubule Network Formation

Next, to investigate the roles of MT network in the regulation of mitophagy in C2C12 myocytes under the oxidative stress, the MT network of C2C12 cells were stained by SiR-tubulin (Fig.4). C2C12 cells under the oxidative stress showed disturbed MT network formation as compared to normal cells, suggesting that the molecular target of oxidative stress likely involves pathways related to MT formation. In line with this finding, it has been reported that trehalose augments autophagy/mitophagy by mTOR-independent pathway [27], but its precise target has not been fully investigated. As shown above, trehalose improves the disturbed maturation process apparently through working on the MT network. The effect Trehalose treatment on MT was thus monitored using SiR-tubulin. Treatment by trehalose of cells exposed to oxidative stress ameliorated the disturbed MT network, suggesting that trehalose can work on maintaining the healthy MT network. To examine whether the signal amelioration of MTs by SiR under trehalose treatment was not due to artifact of measurement, C2C12 cells with trehalose were treated by colchicine. Treatment by colchicine abolished, further supporting that trehalose indeed helps maintain the MT network under oxidative stress condition.

**Figure 4.**
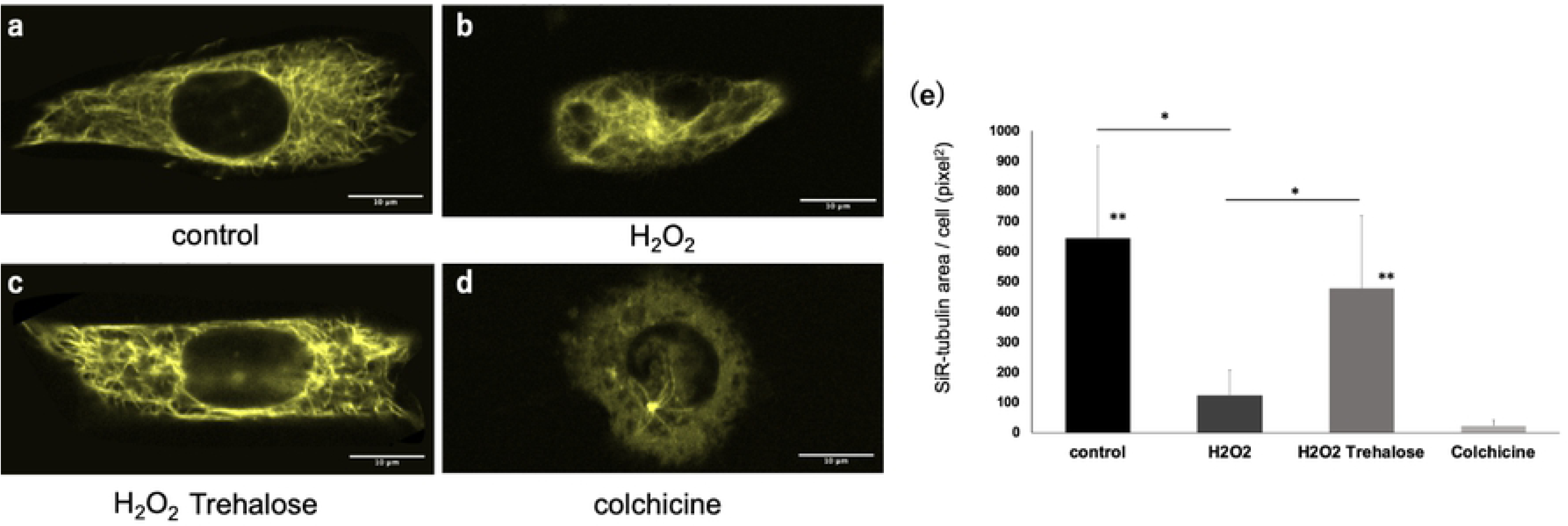
Microtubule network formation is disturbed under oxidative stress. (Top) (a-d) Confocal microscopic images of C2C12 cell stained by SiR-tubulin are shown for the potential effect of H2O2, trehalose and colchicine on the microtubule network. (a) Control group, (b) H_2_O_2_ group, (c) H_2_O_2_ +trehalose group, and (d) H_2_O_2_ +trehalose+colchicine group. Scale bar =10μm. (e) The area of SiR-tubulin was analyzed by ImageJ and shown as the average value of area with standard deviation (N=7). *:p<0.05, **:p<0.05 vs colchicine group by ANOVA with Tukey’s comparison.

### The Effect of Oxidative Stress on the Synthesis of Microtubule

To investigate the mechanism of the beneficial function of trehalose on MTs, the synthesis extent and speed of microtubule was monitored by time-lapse confocal microscopy, using EB1-GFP, the plus-end tracking reporter (Fig.5, Supplementary Video 2). In cells under the subacute oxidative stress, the movement and the number of EB1 significantly decreased as compared to control cells. When cells are treated with trehalose, the diminished movement and number of EB1 was significantly increased in cells under the subacute oxidative stress. These results suggest that trehalose normalizes MT network in cells under oxidative stress conditions either by augmenting the MT synthesis or by inhibiting the MT degradation and thus secondarily affecting the synthesis.

**Figure 5.**
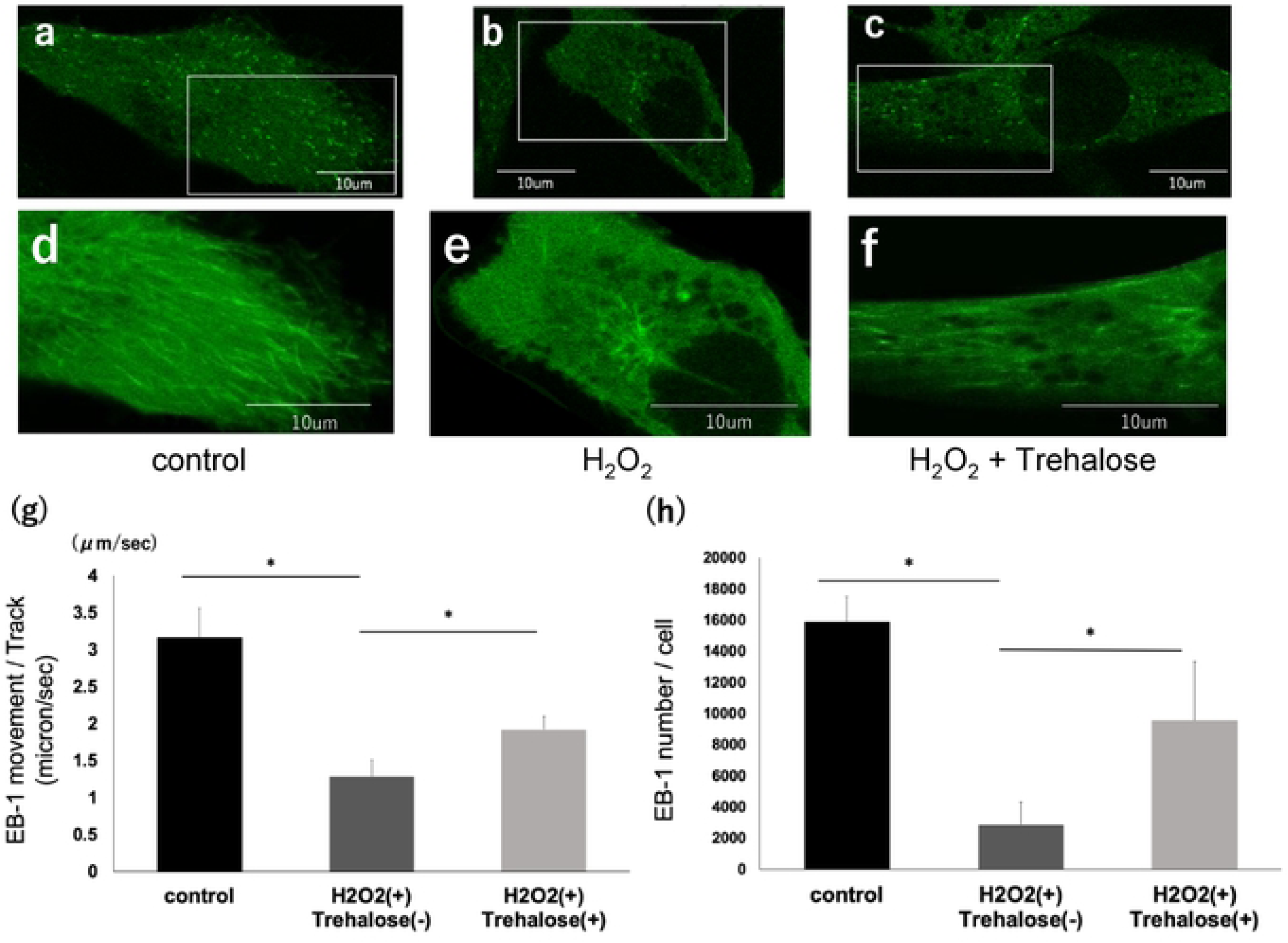
The synthesis ability of microtubule is diminished by oxidative stress. (TOP) (a-c) Confocal microscopic images of C2C12 cell expressing EB1-GFP are shown to analyze the effect of H_2_O_2_ and trehalose on extent and speed of MT synthesis. (a) Control group, (b) H_2_O_2_ group, (c) H_2_O_2_ +trehalose group. Scale bar =10μm. (d-f) Magnified images of the boxed areas in the top (a-c) shown. Synthesized overlay images of ten stacked images of time-lapse observation are shown to document the trajectory of EB1 motion. (d) Control group, (e) H_2_O_2_ group, (f) H_2_O_2_ +trehalose group. Scale bar =10μm. (g, h) The movement speed (g) and number (h) of EB-1 analyzed by ImageJ and shown as the average value of area with standard deviation (N=5). *:p<0.05, **:p<0.05 vs colchicine group by ANOVA with Tukey’s comparison. For video information, refer to Supplementary Video 2. Supplementary Video 2: Time-lapse confocal video image of C2C12 myocytes expressing GFP-EB1 for measuring MT synthesis. Cells are incubated in the presence or absence of H_2_O_2_ with or without trehalose treatment. In controls (H_2_O_2_ -, trehalose -, far left), GFP-EB1 dots, representing the plus-end of elongating MTs, show vigorous movement with greater majority of dots trafficking in the efferent direction. In H_2_O_2_ treated cells without treatment (H_2_O_2_ +, trehalose -, middle), both the numbers and the speed of GFP-EB1 motion are markedly decreased. Trehalose treatment (H_2_O_2_ +, trehalose +, far right) partially rescues the EB1 trafficking.

### Signaling Mechanisms Involved in Disturbed Mitophagy Response

Previous lines of evidence suggested the involvement of Akt/GSK3β signaling in CCCP-induced mitophagy [31] and in the elongation signals of MTs via EB1 activation [32]. Activated Akt functions through the phosphorylation and inhibition of Glycogen Synthase Kinase-3β (GSK3β) [33] leading to the activation of EB1-mediated MT synthesis through CLAPS2 recruitment. We first analyzed the time course of Akt/GSK3β phosphorylation in normal C2C12 cells in response to mitophagy-inducing stimulation (CCCP 5μM and 12.5μM) using Western blot with antibodies against phosphorylated form of Akt (Ser473 p-Akt) and GSK3β (Ser9 p-GSK3β). (Fig.6-a). Both 5μM and 12.5μM stimulation caused rapid phosphorylation of Akt and GSK3β peaking at 15 minutes of stimulation. Since there was no essential difference between 5μM and 12.5μM stimulation, 5μM stimulation was used thereafter unless otherwise stated. Next, to investigate the effect of oxidative stress on the activation of Akt/GSK3β pathway, phosphorylation status of Akt after CCCP stimulation was monitored. Under the oxidative stress, the level of p-Akt and p-GSK3β was significantly decreased as compared to control cells (Fig.6-b, 0.82±0.39 vs. 1.6±0.26; p<0.05; n=5, and 0.70±0.03 vs. 1.17±0.06; p<0.05; n=5, for p-Akt and p-GSK3β, respectively). Trehalose treatment to cells exposed to oxidative stress rescued the deficient signal response of Akt/GSK3β in response to CCCP, as compared to that in cells without treatment (Fig.6-c).

**Figure 6.**
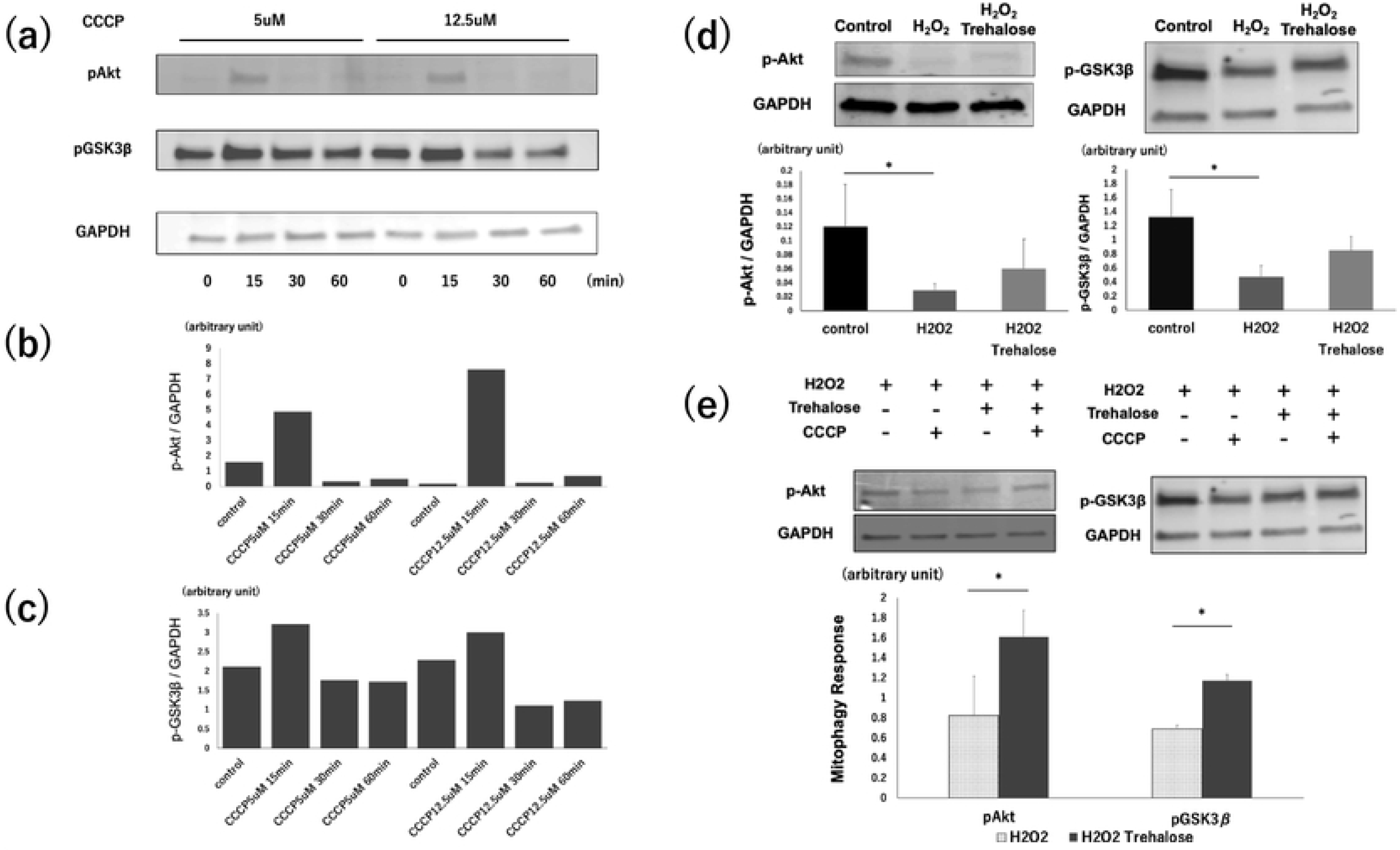
(a-c) Western blot analysis for phospho-Akt (p-Akt) and phospho-GSK3β(p-GSK3β) in the C2C12 myocyte after carbonyl cyanide m-chlorophenyl hydrazone (CCCP) stimulation. In both p-Akt and p-GSK3β expression, CCCP treatment induced rapid phosphorylation of Akt and GSK3β, peaking at 15min. (d) Western blot analysis for p-Akt and p-GSK3β in the C2C12 myocyte with or without H_2_O_2_, trehalose. The p-Akt and p-GSK3β expression significantly decreased in C2C12 cells under the oxidative stress as compared to the control group (n=5, * p<0.05). Trehalose treatment partially reversed the decreased p-Akt and p-GSK3β expression. The results of the GAPDH analysis are shown as an internal control. (e) Western blot analysis for mitophagy response (the increase rate before and after CCCP) in the C2C12 myocyte under the oxidative stress with or without trehalose treatment. Trehalose treatment significantly ameliorated the disturbed p-Akt and p-GSK3β expression (n=5, * p<0.05) under the oxidative stress. The results of the GAPDH analysis are shown as an internal control.

### Superoxide Production From Mitochondria

Mitophagy is one of the major quality control systems of mitochondria that removes defective mitochondria and thus prevents ROS release from the damaged mitochondria. In line with this notion, the superoxide production from mitochondria in C2C12 myocyte was analyzed as an outcome of poor mitophagy response under subacute oxidative stress using a staining dye for mitochondria-derived superoxide, MitoSOX (Fig.7). The ratio of C2C12 with elevated production of superoxide was significantly increased. Trehalose treatment significantly ameliorated the increase of superoxide production, supporting the hypothesis that amelioration of defective mitophagy by trehalose rescues the mitochondria and helps the quality control of mitochondria.

**Figure 7.**
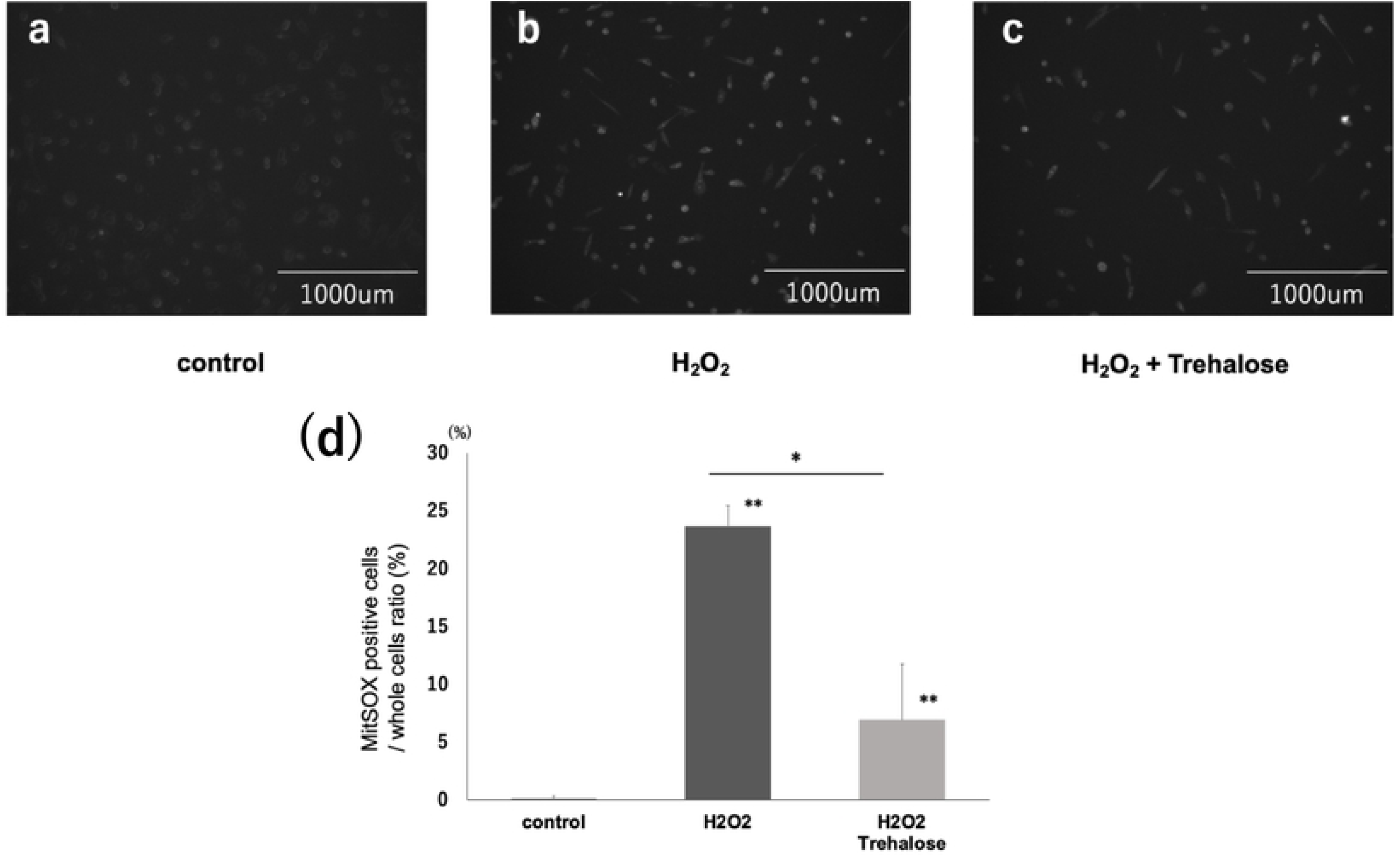
Superoxide production from mitochondria. (a-c) Fluorescence microscopic images of C2C12 cells stained MitoSOX are shown to document the superoxide production from mitochondria after CCCP stimulation. (a) Control group, (b) H_2_O_2_ group, and (c) H_2_O_2_ +trehalose group. Scale bar=1000μm. (d) The ratio of C2C12 cells positively stained by MitoSOX was analyzed by ImageJ and shown as the average value of area with standard deviation (N=6). H_2_O_2_ subacute stimulation increased superoxide production from mitochondria, and H_2_O_2_ +trehalose ameliorated after CCCP administration. *:p<0.05, **:p<0.05 vs control by ANOVA with Tukey’s comparison.

## Discussion

### ROS and Mitophagy Response Defect

In this study, we demonstrated that prolonged oxidative stress decreases the response of mitophagy, especially at the maturation stage, possibly due to the perturbed MT network formation. This is among the first detailed systematic study that analyzed the mechanisms of disturbed mitophagy response, not the basal level of mitophagy, in the cells exposed to an oxidative stress. Furthermore, it was demonstrated that trehalose, a mTOR-independent auto/mitophagy modulator, improves defective mitophagy response in myocytes under the oxidative stress, ameliorating the inhibition of mitophagosomes maturation, presumably by normalizing MT network formation and increasing the synthetic ability of MT. The current study opens a possibility for therapeutic approaches for targeting maturation defect in mitophagy response and for modulating defective MT network formation in treating diseases arising from prolonged oxidative stresses.

### Oxidative Stress and Autophagy/mitophagy - Upregulation or Downregulation?

Although the relationship between ROS and auto/mitophagy has been studied previously [16], most of the previous investigations focused on the mechanisms of ROS-induced upregulation of auto/mitophagy. In these previous studies, the basal levels of auto/mitophagy were analyzed and few studies focused on their response. Furthermore, in many of previous studies, cells or tissues were acutely stimulated for the analyses. Upregulation of auto/mitophagy in such acutely stressed cells is reasonable given that auto/mitophagy evolved as adaptive responses to the external and internal stresses. Importantly, the current finding about the defective mitophagy response is not contradictory to the previous findings for upregulated auto/mitophagy under oxidative stress. One has to take into account whether the analyses are performed to quantify the basal level of auto/mitophagy, or the changes in their responses of auto/mitophagy to stimulations. In biological systems, these two often show opposite trends. For example, in type 2 diabetes mellitus (T2DM), despite the fact that pathophysiology of many disease symptoms can be explained by insulin resistance, or the defective response of intracellular signals to insulin stimulation in many organs, even though the basal level of plasma insulin is often elevated in many patients. Despite the serum insulin elevation, suppression or blockade of insulin signal would not be considered a reasonable therapeutic approach. Likewise, although many previous studies reported upregulation of the basal level of auto/mitophagy, blocking autophagy or mitophagy in oxidative stress related diseases may not provide a useful therapeutic approach, especially given that auto/mitophagy are mostly cell protective response in many situations.

### Mitophagy Defect and Human Diseases Related to Oxidative Stress

Mitophagy, a form of macroautophagy that selectively degrades damaged mitochondria, is considered as one of the major quality control system of mitochondria, and helps maintaining the integrity and the functions of healthy mitochondria [34]. Mitophagy defect is currently assumed to be an essential pathogenic factor in cancer, neurodegeneration, metabolic disorders, muscle atrophy, inflammation and organ failure caused by sepsis [35, 36].

Among many other diseases, prolonged increase of oxidative stress is considered one of the key mechanisms in sepsis or burn injury induced organ dysfunctions. In our experiments, subacute H_2_O_2_ stimulation significantly inhibited mitophagy response, after CCCP stimulation (Fig.1,2) blocked mitophagosomes maturation (Fig.3), resulting in elevated superoxide generation from the defective mitochondria (Fig.7). These data suggest that subacute oxidative stress leads to mitophagy defect in myocyte after CCCP stimulation. Trehalose, however, improved the defects in maturation under the subacute oxidative stress (Fig.3) and thus will be a promising drug candidate for treating the mitophagy defect.

### Microtubule and Mitophagy Maturation

MT forms a dynamic cytoskeletal structure. Both integrity of healthy MT and the normal functions of MT-based motors have been shown essential in autophagy [22]. The centripetal movement of mature autophagosomes prior to fusion with lysosomes requires stable MT [37]. A key mechanism of auto/mitophagosome flux is the transport of auto/mitophagosomes along MTs toward lysosomes to form mature autophagolysosomes (autolysosomes) [38]. In this study, observation C2C12 cells stained by SiR-tubulin showed that the subacute oxidative stress abolished MT network formation as compared to normal cells (Fig4). Observation of tfLC3-expressing cells demonstrated that the subacute oxidative stress inhibited mitophagosomes maturation (Fig3), implying that the unstable MT could be one of the important mechanisms of mitophagy defect under the subacute oxidative stress. This notion is supported by the finding that trehalose treatment ameliorated MT defect and mitigated mitophagy maturation defect but that colchicine abolished the beneficial effect of trehalose. These data suggest that MT serves as an essential guide rail for mitophagosome vesicle trafficking and thus are a key component of mitophagy pathway involved in disease conditions

Phosphoinositide-3-Kinase (PI3-K) and the downstream kinases Akt and GSK3β have recently been implicated in regulating both MT dynamics and organization [39]. Akt and GSK3β coordinately regulate the phosphorylation status of MT end capping molecules including CLASP2 and EB1, thus regulating their recruitment to the synthesis plus end of MTs[40]. Importantly, auto/mitophagy stimulating signal activate this pathway [31, 41]. The results of Western Blot analysis showed that the subacute oxidative stress significantly decreased the pAkt and GSK3β expression as compared to control cells (Fig.6-b, c), suggesting that subacute oxidative stress can affect MT synthesis by invoking signal resistance in PI3K/Akt/GSK3β pathway.

### Trehalose Function as a Modulator of Mitophagy

It has been established that trehalose is a modulator of auto/mitophagy pathway, augmenting these stress adaptation functions by mTOR-independent mechanisms in COS-7 cells and mouse embryonic fibroblasts (MEFs) [42, 43]. Trehalose also upregulates p62 expression and activates the autophagy flux in Hepa1-6 cells and MEFs [9]. In our study, trehalose can improve the synthesis ability of MT and thus ameliorates the defective MT network under the subacute oxidative stress (Fig.4). Thus repaired MT network improved the oxidative stress-induced inhibition of mitophagosomes maturation in trehalose treated cells. Notably, colchicine, a drug that destabilizes MT, abolished the beneficial effect of trehalose on mitophagy (Fig.3&4). In the detailed analyses of the signal transduction mechanisms, trehalose treatment significantly increased the disturbed Akt/GSK3β signaling in cells under the oxidative stress (Fig.6-b, c). Many therapeutic approaches have been previously suggested in human diseases or their murine equivalent models where auto/mitophagy are defective. In many of previous studies, one of the proposed strategies was to boost auto/mitophagy. The current study suggests, however, that auto/mitophagy defect in diseases can involve maturation defect, or the blocked flux. To forcefully stimulate the upstream activation of auto/mitophagy in such condition where the downstream is blocked, (e.g. by disturbed MT network as shown in the current study), may not serve an effective therapeutic approach. Thus, the importance of finding the precise target of auto/mitophagy defect and ameliorating the perturbation, cannot be overemphasized.

Oxidative stress-induced MT disturbance can be a pivotal event in the critical illness-related mitochondrial dysfunction in skeletal muscles, and needs further investigation. Trehalose can ameliorate the perturbed MTs by increasing the synthetic ability of MT, and normalizing the disturbed MTs can serve a novel therapeutic agent in critical illnesses.

## Conclusions

Oxidative stress decreases the response of mitophagy and abolishes MT network formation. Trehalose improves the synthetic ability of MT by increasing the number and movement of the MT plus end-capping molecule, EB1, suggesting that trehalose augments MT network synthesis. Observation of tfLC3-expressing cells demonstrates that trehalose also improve the inhibition of mitophagosomes maturation under the oxidative stress. Finally, trehalose normalizes MT network formation by increasing the synthetic ability of MT. All the obtained data suggest that normalizing the disturbed MTs under oxidative stress can prevent the mitophagy dysfunction under the oxidative stress and serve a novel therapeutic target.

## Acknowledgements

We used Shiners Morphology Core and Simches PMB Microscope Core for acquiring microscopy data, Shriners Genomics and Proteomics Core for performing biochemical analyses. We thank Daniel Fong for administrative assistance.

## Reference

1. Ueki R, Liu L, Kashiwagi S, Kaneki M, Khan MA, Hirose M, Tompkins RG, Martyn JA, Yasuhara S (2016) Role of Elevated Fibrinogen in Burn-Induced Mitochondrial Dysfunction: Protective Effects of Glycyrrhizin. Shock 46: 382–389.

2. Batt J, Herridge M, Dos Santos C (2017) Mechanism of ICU-acquired weakness: skeletal muscle loss in critical illness. Intensive Care Med 43: 1844–1846.

3. Kraft R, Herndon DN, Finnerty CC, Shahrokhi S, Jeschke MG (2014) Occurrence of multiorgan dysfunction in pediatric burn patients: incidence and clinical outcome. Ann Surg 259: 381–387.

4. Warren DK, Shukla SJ, Olsen MA, Kollef MH, Hollenbeak CS, Cox MJ, Cohen MM, Fraser VJ (2003) Outcome and attributable cost of ventilator-associated pneumonia among intensive care unit patients in a suburban medical center. Crit Care Med 31: 1312–1317.

5. Hermans G, Van den Berghe G (2015) Clinical review: intensive care unit acquired weakness. Crit Care 19: 274.

6. Winkelman C (2010) The role of inflammation in ICU-acquired weakness. Crit Care 14: 186.

7. Romanello V, Sandri M (2015) Mitochondrial Quality Control and Muscle Mass Maintenance. Front Physiol 6: 422.

8. Ohsumi Y (2014) Historical landmarks of autophagy research. Cell Res 24: 9–23.

9. Mizunoe Y, Kobayashi M, Sudo Y, Watanabe S, Yasukawa H, Natori D, Hoshino A, Negishi A, Okita N, Komatsu M, Higami Y (2018) Trehalose protects against oxidative stress by regulating the Keap1-Nrf2 and autophagy pathways. Redox Biol 15: 115–124.

10. Hosokawa S, Koseki H, Nagashima M, Maeyama Y, Yomogida K, Mehr C, Rutledge M, Greenfeld H, Kaneki M, Tompkins RG, Martyn JA, Yasuhara SE (2013) Title efficacy of phosphodiesterase 5 inhibitor on distant burn-induced muscle autophagy, microcirculation, and survival rate. Am J Physiol Endocrinol Metab 304: E922–933.

11. Masiero E, Agatea L, Mammucari C, Blaauw B, Loro E, Komatsu M, Metzger D, Reggiani C, Schiaffino S, Sandri M (2009) Autophagy is required to maintain muscle mass. Cell Metab 10: 507–515.

12. Jiroutkova K, Krajcova A, Ziak J, Fric M, Waldauf P, Dzupa V, Gojda J, Nemcova-Furstova V, Kovar J, Elkalaf M, Trnka J, Duska F (2015) Mitochondrial function in skeletal muscle of patients with protracted critical illness and ICU-acquired weakness. Crit Care 19: 448.

13. Filomeni G, Desideri E, Cardaci S, Rotilio G, Ciriolo MR (2010) Under the ROS…thiol network is the principal suspect for autophagy commitment. Autophagy 6: 999–1005.

14. Scherz-Shouval R, Elazar Z (2007) ROS, mitochondria and the regulation of autophagy. Trends Cell Biol 17: 422–427.

15. Morishita H, Mizushima N (2019) Diverse Cellular Roles of Autophagy. Annu Rev Cell Dev Biol 35: 453–475.

16. Filomeni G, De Zio D, Cecconi F (2015) Oxidative stress and autophagy: the clash between damage and metabolic needs. Cell Death Differ 22: 377–388.

17. Alberti C, Brun-Buisson C, Chevret S, Antonelli M, Goodman SV, Martin C, Moreno R, Ochagavia AR, Palazzo M, Werdan K, Le Gall JR, European Sepsis Study G (2005) Systemic inflammatory response and progression to severe sepsis in critically ill infected patients. Am J Respir Crit Care Med 171: 461–468.

18. Skrupky LP, Kerby PW, Hotchkiss RS (2011) Advances in the management of sepsis and the understanding of key immunologic defects. Anesthesiology 115: 1349–1362.

19. Ueki R, Kashiwagi A, Hirose M, Yu YM, Martyn JA, Yasuhara S (2015) Mitophagy resistance in skeletal muscles after burn injury. ABA Meeting.

20. Yasuda N, Shakuo T, Kashiwagi A, Tamura T, Kaneki M, Khan M, Martyn JA, Yasuhara S (2019) Evidence for burn-induced perturbation in mitophagic response (mitophagy resistance) based on skeletal muscle cell culture experiments. American Burn Association Annual Meeting.

21. Scherz-Shouval R, Shvets E, Elazar Z (2007) Oxidation as a post-translational modification that regulates autophagy. Autophagy 3: 371–373.

22. Mackeh R, Perdiz D, Lorin S, Codogno P, Pous C (2013) Autophagy and microtubules - new story, old players. J Cell Sci 126: 1071–1080.

23. Hengherr S, Heyer AG, Kohler HR, Schill RO (2008) Trehalose and anhydrobiosis in tardigrades--evidence for divergence in responses to dehydration. FEBS J 275: 281–288.

24. Bradbury J (2001) Of tardigrades, trehalose, and tissue engineering. Lancet 358: 392.

25. Campbell LH, Brockbank KG (2012) Culturing with trehalose produces viable endothelial cells after cryopreservation. Cryobiology 64: 240–244.

26. Eroglu A, Bailey SE, Toner M, Toth TL (2009) Successful cryopreservation of mouse oocytes by using low concentrations of trehalose and dimethylsulfoxide. Biol Reprod 80: 70–78.

27. Sarkar S, Davies JE, Huang Z, Tunnacliffe A, Rubinsztein DC (2007) Trehalose, a novel mTOR-independent autophagy enhancer, accelerates the clearance of mutant huntingtin and alpha-synuclein. J Biol Chem 282: 5641–5652.

28. Tanaka M, Machida Y, Niu S, Ikeda T, Jana NR, Doi H, Kurosawa M, Nekooki M, Nukina N (2004) Trehalose alleviates polyglutamine-mediated pathology in a mouse model of Huntington disease. Nat Med 10: 148–154.

29. Draberova E, Sulimenko V, Sulimenko T, Bohm KJ, Draber P (2010) Recovery of tubulin functions after freeze-drying in the presence of trehalose. Anal Biochem 397: 67–72.

30. Sadhu DN, Lundberg MS, Burghardt RC, Ramos KS (1994) c-Ha-rasEJ transfection of rat aortic smooth muscle cells induces epidermal growth factor responsiveness and characteristics of a malignant phenotype. J Cell Physiol 161: 490–500.

31. Soutar MPM, Kempthorne L, Miyakawa S, Annuario E, Melandri D, Harley J, O’Sullivan GA, Wray S, Hancock DC, Cookson MR, Downward J, Carlton M, Plun-Favreau H (2018) AKT signalling selectively regulates PINK1 mitophagy in SHSY5Y cells and human iPSC-derived neurons. Sci Rep 8: 8855.

32. Schmidt N, Basu S, Sladecek S, Gatti S, van Haren J, Treves S, Pielage J, Galjart N, Brenner HR (2012) Agrin regulates CLASP2-mediated capture of microtubules at the neuromuscular junction synaptic membrane. J Cell Biol 198: 421–437.

33. Cross DA, Alessi DR, Cohen P, Andjelkovich M, Hemmings BA (1995) Inhibition of glycogen synthase kinase-3 by insulin mediated by protein kinase B. Nature 378: 785–789.

34. Yin X, Xin H, Mao S, Wu G, Guo L (2019) The Role of Autophagy in Sepsis: Protection and Injury to Organs. Front Physiol 10: 1071.

35. Springer MZ, Macleod KF (2016) In Brief: Mitophagy: mechanisms and role in human disease. J Pathol 240: 253–255.

36. Thiessen SE, Derese I, Derde S, Dufour T, Pauwels L, Bekhuis Y, Pintelon I, Martinet W, Van den Berghe G, Vanhorebeek I (2017) The Role of Autophagy in Critical Illness-induced Liver Damage. Sci Rep 7: 14150.

37. Kast DJ, Dominguez R (2017) The Cytoskeleton-Autophagy Connection. Curr Biol 27: R318–R326.

38. Geeraert C, Ratier A, Pfisterer SG, Perdiz D, Cantaloube I, Rouault A, Pattingre S, Proikas-Cezanne T, Codogno P, Pous C (2010) Starvation-induced hyperacetylation of tubulin is required for the stimulation of autophagy by nutrient deprivation. J Biol Chem 285: 24184–24194.

39. Nehlig A, Molina A, Rodrigues-Ferreira S, Honore S, Nahmias C (2017) Regulation of end-binding protein EB1 in the control of microtubule dynamics. Cell Mol Life Sci 74: 2381–2393.

40. Basu S, Sladecek S, Pemble H, Wittmann T, Slotman JA, van Cappellen W, Brenner HR, Galjart N (2014) Acetylcholine receptor (AChR) clustering is regulated both by glycogen synthase kinase 3beta (GSK3beta)-dependent phosphorylation and the level of CLIP-associated protein 2 (CLASP2) mediating the capture of microtubule plus-ends. J Biol Chem 289: 30857–30867.

41. Shi Y, Yan H, Frost P, Gera J, Lichtenstein A (2005) Mammalian target of rapamycin inhibitors activate the AKT kinase in multiple myeloma cells by up-regulating the insulin-like growth factor receptor/insulin receptor substrate-1/phosphatidylinositol 3-kinase cascade. Mol Cancer Ther 4: 1533–1540.

42. Fleming A, Noda T, Yoshimori T, Rubinsztein DC (2011) Chemical modulators of autophagy as biological probes and potential therapeutics. Nat Chem Biol 7: 9–17.

43. Sarkar S (2013) Regulation of autophagy by mTOR-dependent and mTOR-independent pathways: autophagy dysfunction in neurodegenerative diseases and therapeutic application of autophagy enhancers. Biochem Soc Trans 41: 1103–1130.

